# Identification and characterization of bifunctional pyrophosphate-fructose-6-phosphate 1-phosphotransferase from citrus greening disease-causing α-proteobacterium

**DOI:** 10.1101/2025.04.04.647159

**Authors:** Sandra Kathott Prakash, Etisha Arun, Khushhal Rathore, Mrugendra Gubyad, Dilip Kumar Ghosh, Ashwani Kumar Sharma, Jitin Singla

## Abstract

*Candidatus* Liberibacter asiaticus (*C*Las) is an obligate intracellular pathogen that causes citrus greening disease. Prolonged host association has driven reductive evolution and functional adaptation, leaving central carbon metabolism disrupted by the absence of phosphoglucose isomerase (*pgi*) in glycolysis and transaldolase in the pentose phosphate pathway (PPP). Here, we characterized the bifunctional enzyme, pyrophosphate-fructose-6-phosphate 1-phosphotransferase (PFP), that connects glycolysis and PPP. PFP primarily converts D-fructose 6-phosphate (F6P) to D-fructose 1,6-bisphosphate (FBP). The proposed alternative function of PFP is the conversion of D-sedoheptulose 7-phosphate (S7P) to D-sedoheptulose 1,7-bisphosphate (SBP), and this reaction is part of the sedoheptulose 1,7-bisphosphate pathway (SBPP). To validate this, the recombinant *C*Las PFP was heterologously expressed, purified, and biochemically characterized. The steady-state kinetic studies revealed *C*Las PFP showed substrate affinity towards both substrates, F6P and S7P, with Km values of 67 ± 5 μM and 58 ± 3 μM, respectively. In silico analysis revealed distinct motifs that reflect specificity for phosphoryl donors among phosphofructokinase (PFK) isoforms and showed that PFP may be ancestral. Molecular docking and simulation studies of PFP-complexes with F6P and S7P showed stable binding and interaction. Biophysical characterization showed that PFP loses conformational stability above 50 °C and at pH 11.0. This is the first report of characterization of PFP from CLas, demonstrating that the protein is a functional, structurally stable enzyme with dual substrate specificity.

## 1. Introduction

Obligate intracellular microorganisms rely on the host for metabolism and replication while evading host defenses^[1]^. Prolonged cohabitation of a microorganism within a host led to reduced population size and lower genetic diversity^[2]^. The transition from a free-living to a host-restricted lifestyle alters or eliminates essential genes, a process termed reductive evolution, with consequent loss of metabolic functions^[1,3]^. Some notable examples of obligate intracellular bacterial species include the genera *Chlamydia, Rickettsia, Buchnera,* and *Mycoplasma^[4–7]^*. Genome and proteome analyses of such bacteria reveal pathways that are incomplete due to missing genes, making them dependent on host metabolites for survival^[7]^. Central carbon metabolism is frequently affected, and disrupted glycolysis may be compensated by other pathways, such as the pentose phosphate pathway and the Entner-Doudoroff pathway^[8]^. Where these compensatory pathways are also disrupted, alternative routes have emerged through the evolution of new genes or neofunctionalization.

*Candidatus* Liberibacter asiaticus (*C*Las) is a gram-negative, phloem-limited alphaproteobacterium transmitted by the insect vector *Diaphorina citri*, commonly known as *psyllid*, causing the disease called “citrus greening” or Huanglongbing (HLB) in *citrus spp^[9]^*. The close association of CLas with the hosts (*citrus* and *psyllid*) has led to reduced metabolic capability^[10]^, hindering the efforts to establish an axenic culture of CLas^[11–13]^. Although individual CLas proteins have been studied, functional characterization, metabolomics, and genetic manipulation remain limited due to the inability to grow the bacterium in a host-free system. *Liberibacter crescens* (Lcr) strain BT-1 was the first culturable bacterium of the genus *Liberibacter* isolated from infected leaf sap of Babaco Mountain papaya (*Carica stipulata* × *C. pubescens*)^[14]^. Lcr BT-1 is non-pathogenic and does not have any insect or plant host^[15]^.

Previous studies have shown that *C*Las lacks essential genes associated with the glycolysis pathway, including phosphoglucose isomerase (*pgi*), and this reaction can be compensated via the pentose phosphate pathway (PPP), but this possibility is compromised due to the absence of transaldolase^[16]^. Comparative genomics of the close relative *Candidatus* Liberibacter solanacearum (*C*Lso) revealed loss of phosphoenolpyruvate-phosphotransferase system (PTS), a partial PPP, and absence of energy metabolism enzymes, including polyphosphate kinase, inorganic diphosphatase, terminal oxidases: cytochrome c oxidase and cytochrome bd complex^[17]^. It has been reported that *C*Las has an incomplete methylglyoxal detoxification system^[18]^. Also, *C*Las encodes an ATP translocase (GU011685.1) to import ATP from the host and meet the energy requirement^[19]^.

In the present study, we carried out comparative genome analysis of *C*Las to identify non-canonical reactions catalyzed by enzymes involved in carbon metabolism. The metabolic modeling revealed an intermediate reaction involving the pyrophosphate-fructose-6-phosphate 1-phosphotransferase (PFP) in the pentose phosphate pathway, beyond its usual role in glycolysis. This reaction is part of the sedoheptulose 1,7-bisphosphate pathway (SBPP), a variant of the PPP, in which PFP, along with fructose-bisphosphate aldolase (FBPA), is postulated to bypass the function of transaldolase in some organisms. The potential role of *C*Las PFP was investigated further using *in silico* and *in vitro* approaches. The sequence-and structure-based analysis of *C*Las PFP and its homologs was conducted to explore the evolutionary adaptation of PFP across species. Further, we report the expression, purification, and biochemical and biophysical characterization of *C*Las PFP to validate this multifunctional role and structural stability.

## 2. Results

### 2.1 Genome analysis and Functional annotation

The genome annotation of *C*Las gxpsy and Lcr BT-1 was obtained from RAST annotation server and was used to build metabolic models for comparison. Compared to Lcr BT-1 (∼1.5 Mb, 1,269 genes), *C*Las gxpsy has a significantly reduced genome of ∼1.28 Mb and 1,078 protein-coding genes. Evaluation of the functional annotation results for *C*Las and Lcr BT-1 revealed missing genes and additional genes in the central carbon metabolism of *C*Las (**Table S1**). Comparative sequence analysis with its homologs showed that *C*Las gxpsy has lost critical carbon metabolism genes, including *pgi* and transaldolase **(Table S2).** Instead of 6-phosphofructokinase (PFK-1), both *C*Las gxpsy and Lcr BT-1 encode PFP (EC 2.7.1.90), which catalyzes the conversion of D-fructose 6-phosphate (F6P) to D-fructose 1,6-bisphosphate (FBP) (**Figure 1**). Comparison of genome sequences showed that *Liberibacter spp., Prevotella copri, Clostridium thermosuccinogenes,* and *Entamoeba histolytica* encode PFP protein but lack transaldolase. Also, some species, including *Afipia carboxidovorans, Azorhizobium caulinodans, Bartonella spp., Brucella spp., Methylobacterium radiotolerans, Methylocella silvestris, Methylorubrum extorquens, Rhodomicrobium vannielii,* and *Rhodopseudomonas palustris, lack* PFK-1 but have transaldolase. These results show that the adaptation of central carbon metabolism varies across different species and are tabulated in **Table S2**.

**Figure 1.**
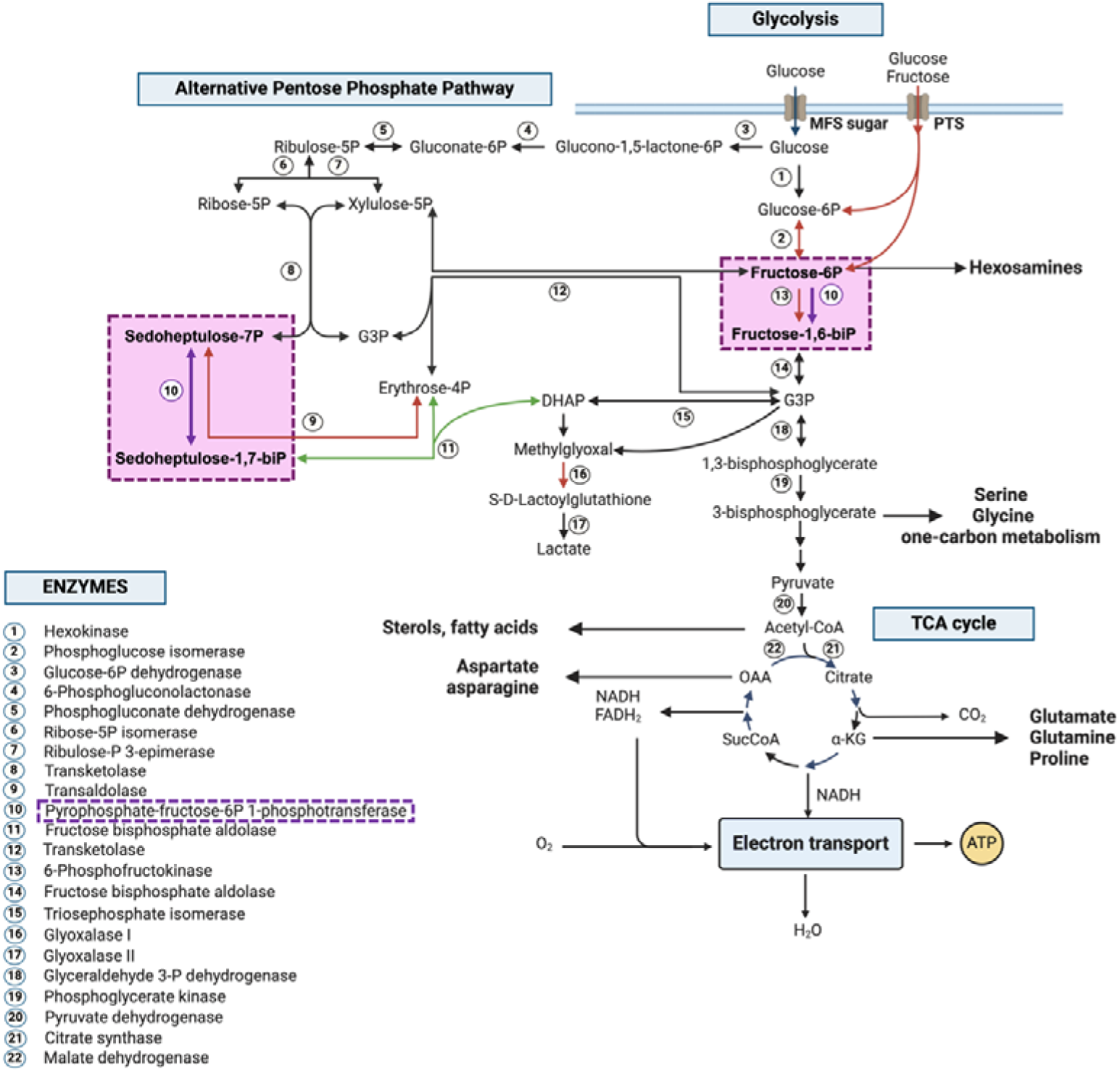
The representation of central carbon metabolism. Genes present in *C*Las gxpsy are indicated by black arrows, while absent genes are marked with red arrows. Alternative reactions catalyzed by PFP and FBPA to compensate for the absence of transaldolase are indicated by the violet and green arrows, respectively. The metabolic pathway is created in BioRender. KP, S. (2025) https://BioRender.com/c31y167

The metabolic model of *C*Las gxpsy built using minimal glucose media on KBase server shows a non-canonical reaction catalyzed by PFP to form D-sedoheptulose 1,7-bisphosphate (SBP) from D-sedoheptulose 7-phosphate (S7P) in PPP as an intermediate reaction (**Figure 1**). This reaction was not identified in Lcr BT-1. A similar pathway involving PFP and FBPA (**Figure 1)** to circumvent the loss of transaldolase has been reported in other organisms^[20–22]^.

### 2.2 Sequence analysis of *C*Las PFP

The sequence similarity search revealed that CLas encodes a single type of phosphofructokinase (PFK) protein, PFP, which belongs to phosphofructokinase type A (PFKA; IPR050929). The bioinformatics tool ProtParam predicted a stable protein with theoretical pI, extinction coefficient, and molecular weight of 6.56, 34755 M^-1^cm^-1^, and 44.4kDa, respectively. Sequences analyzed spanned the bacterial phyla proteobacteria, bacteroidota, actinomycetota, and firmicutes, and the eukaryotic phyla evosea, euglenozoa, and apicomplexa. Several species encode two or three variants of PFK, with motif variations reflecting specificity for different phosphoryl donors (PPi, ATP, and an ATP-dependent variant with an atypical amino acid combination denoted as PFK* and PFK**^#^**). Multiple sequence alignment of PFP sequence from CLas shows variable identity with *Liberibacter spp.* viz *Candidatus* Liberibacter africanus (*C*Laf) (85.2%), *C*Lso (81.5%), *Candidatus* Liberibacter americanus (*C*Lam) (74.5%), and Lcr BT-1 (64.8%). Other homologs included *Cutibacterium acnes* (PFP: 51.7%, PFK-1: 25.2%, PFK*: 29.7%), *E. coli* (PFK-1: 27.9%), *Prevotella copri* (PFP: 28.7%), *E.histolytica* (PFP: 24%, PFK*: 28%), *Clostridium thermosuccinogenes* (PFP: 27%, PFK-1: 24% PFK*: 26%), *Clostridium thermocellum* (PFP: 26%, PFK-1: 24%), *Chlamydia psittaci* (PFP: 23.8%, PFK*: 27.6%), *Chlamydia trachomatis* (PFK*: 26.1%, PFK**^#^**: 27.1%). Despite the variable identities, the alignment indicates a conserved catalytic domain with minimal functional change in substrate binding (**Figure 2**).

**Figure 2.**
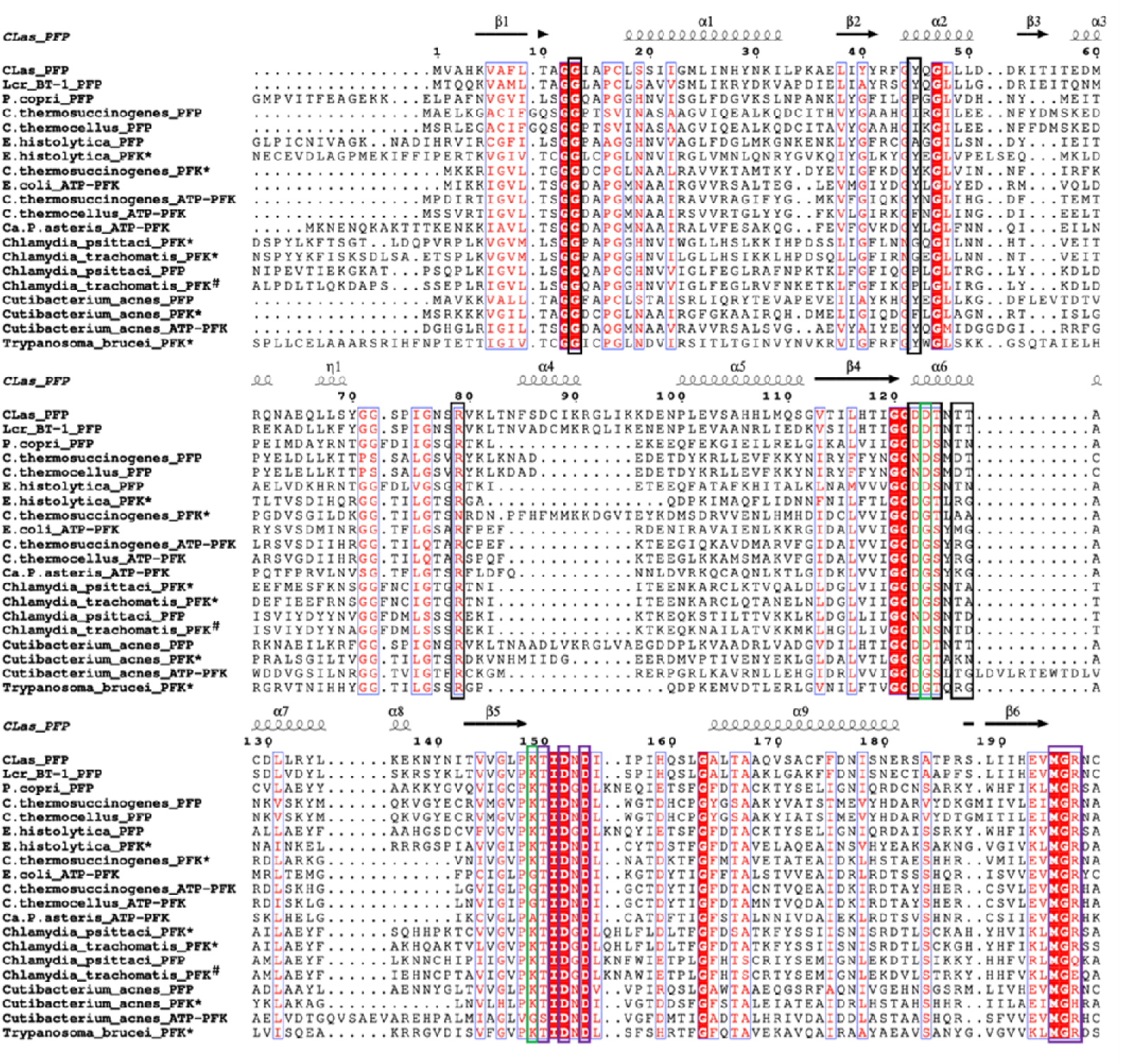
Multiple sequence alignment of phosphofructokinase protein sequences from homologous species. The substrate binding sites are shown in purple boxes, and the PPi binding sites are indicated in black boxes. The green box highlights the atypical amino acid combination observed among different species.

Substrates and co-substrates binding domains were identified using the NCBI conserved domain search tool^[23]^. Conserved motifs were compared across homologs to identify residues influencing ATP/PPi binding. Sequence analysis of PFP (PPi-dependent) proteins revealed two conserved signature motifs: GG(D/N)**D** and **K**TIDXD. The highlighted residues correspond to D123 and K149 in *C*Las PFP (**Figure 2**). Substitutions at these sites were observed in PFK-1 (ATP-dependent), with the patterns GGD**G** and **G**TIDXD. Another variant, PFK*, showed affinity for ATP with motifs GGDG and KTIDXD, but in *Cutibacterium acnes,* motifs were GGG**G** and **K**TIDXD. PFK-1 from *Phytoplasma spp*. carried atypical motifs, GGDG and ATIDXD, with glycine substituted by alanine, unlike the well-conserved PFK-1 and PFK* motifs. PFK# from *Chlamydia trachomatis* showed a signature pattern of GGDN and KTIDXD, also unique among the homologs (**Figure 2**). Other active-site substitutions relative to CLas PFP included Y45 →(P/I/F/G), R79 → (N/K), and T126 → (D/R/K/M). Substrate binding sites are conserved among homologs except at position 197, where arginine is replaced by glutamine, glutamate, and histidine in *Chlamydia psittaci* PFP, *Chlamydia trachomatis* PFK**^#^**, and *Cutibacterium acnes* PFK*, respectively (**Figure 2**).

To infer evolutionary divergence and ancestral points, a maximum-likelihood tree was constructed in MEGA11^[24]^ using 52 nucleotide sequences from gut bacteria, plant pathogens, and protozoan parasites across different phyla; the tree had a maximum log-likelihood of –13705.2 and a genetic distance of 0.5. A mid-point rooted tree showed two major ancestral nodes from which different PFK variants diverged (**Figure 3**). PFK variants were unevenly distributed across species, with two or three encoded in a single genome in some cases (*Clostridium spp., E. histolytica, Cutibacterium acnes, Roseburia intestinalis, Borrelia burgdorferi, Prevotella spp., Leyella stercorea, Bacteroides gallinarum, Buchnera aphidicola, Chlamydia spp., Toxoplasma gondii*), indicating the complex evolutionary pattern of PFK.

**Figure 3.**
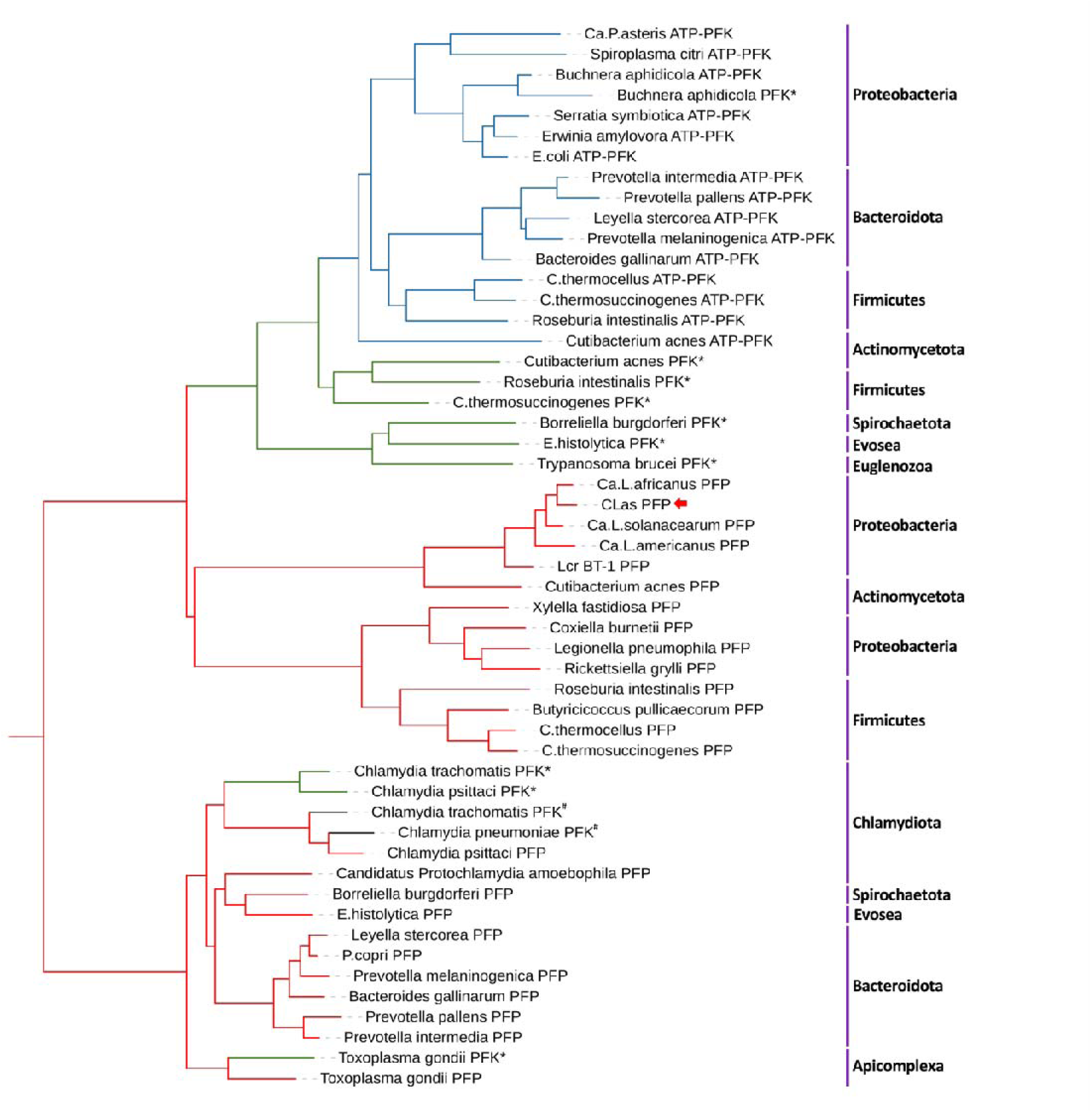
The phylogenetic tree generated using MEGA11. Different phosphofructokinase (PFK) variants are represented with different colors: pyrophosphate-fructose-6-phosphate 1-phosphotransferase (PFP) (red), 6-phosphofructokinase (PFK-1 or ATP-PFK) (blue), ATP-dependent variant of PFK: PFK* (green), PFK# (black).

One of the major ancestral nodes had two clades: one diverging to PFP from proteobacteria, firmicutes, and actinomycetota, and the other to PFK-1 and PFK* (**Figure 3**). *C*Las PFP is closely related to *C*Laf PFP, forming a sister clade, with Lcr BT-1 forming a basal lineage to *Liberibacter spp.* The closest taxa that diverged from the same last common ancestor (LCA) as *Liberibacter spp.* is *Cutibacterium acnes* PFP. PFP proteins sharing the same LCA as *Liberibacter spp*. include those from obligate intracellular bacteria (*Coxiella burnetii*, *Rickettsiella grylli*), xylem-limited *Xylella fastidiosa*, facultative intracellular *Legionella pneumophila*, gut bacteria (*Butyricoccus pullicaecorum*, *Roseburia intestinalis*), and free-living *Clostridium spp.* The tree showed that PFK-1 and PFK* homologs descend from a common node, and this clade shares the same LCA as PFP proteins, indicating an early divergence of PFP.

### 2.3 Molecular modeling of CLas PFP

The structure-based comparison of the PFK variants was carried out to understand the influence of substitutions at the active site on conformation. Among the structurally characterized PFK homologs from various species, *Prometheoarchaeum syntrophicum* (PDB ID:9CIR), *Marinobacter nauticus* (PDB ID:3K2Q), *Nitrosospira multiformis* (PDB ID: 3HNO), and *Borrelia burgdorferi* (PDB ID: 1KZH) have PFP motif, and *Trypanosoma brucei* (PDB ID: 2HIG) has PFK* motif. The three-dimensional structures of PFP predicted with AlphaFold3^[25]^, ESMFold^[26]^, and RoseTTAFold^[27]^ were superimposed on the catalytic domain (**Figure 4a).** 3K2Q, which gave the lowest RMSD among the homologous structures, was used as the reference structure. Comparing 3K2Q with AlphaFold3, ESMFold, and RoseTTAFold structures give RMSD values of 1.54 Å, 1.71 Å, and 2.02 Å, respectively. The AlphaFold3 model was used for further study.

**Figure 4.**
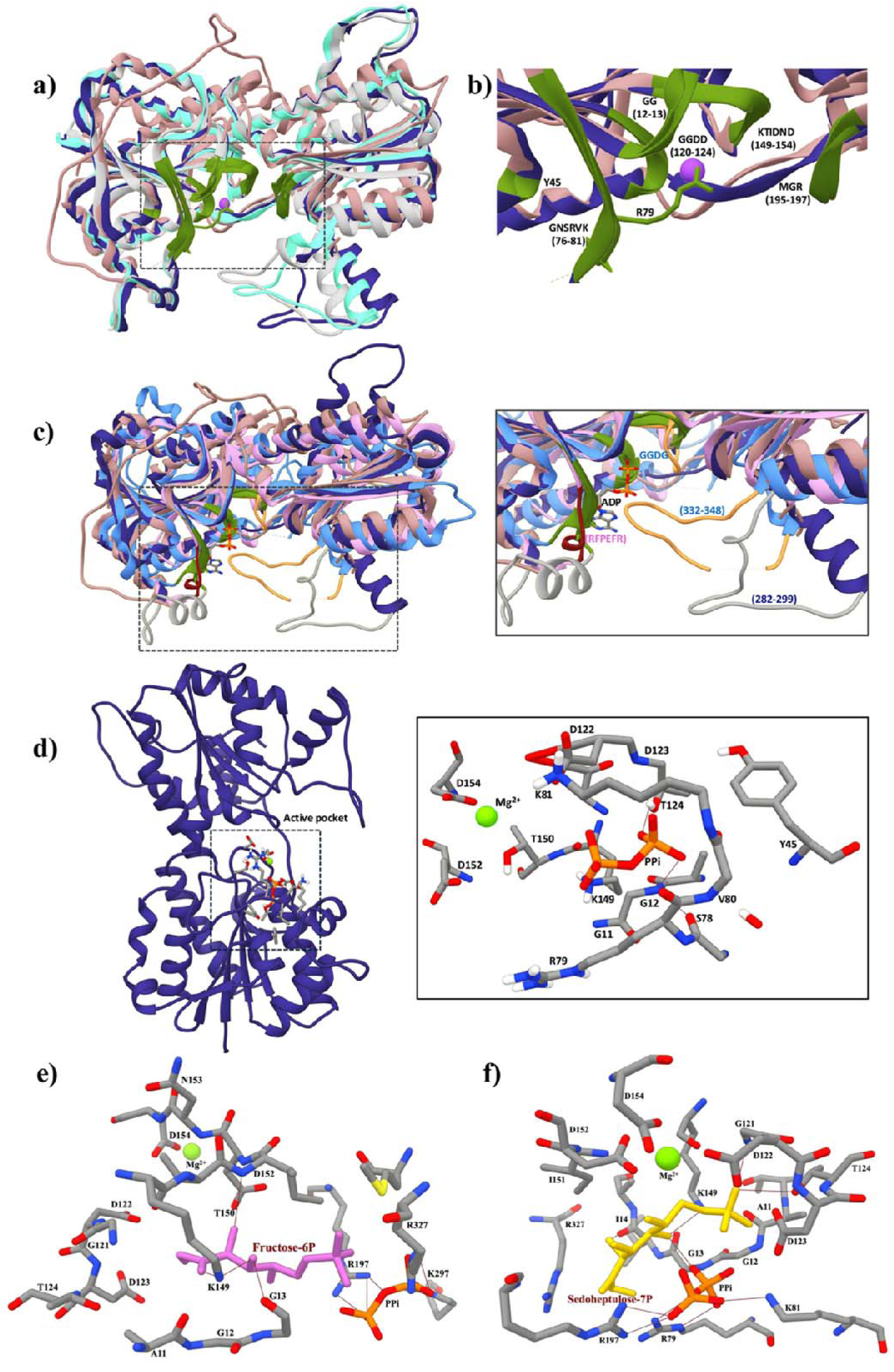
The PFP structure highlights the α/β domain and the catalytic domain **(a)** The superimposed structures of PFP predicted using AlphaFold3 (blue), ESMFold (cyan), RoseTTAFold (light grey), and *Marinobacter nauticus* PFP (3K2Q, light brown) are shown in cartoon representation, and the catalytic site is highlighted in green color. (**b**) structural comparison of 3K2Q with *C*Las PFP model, highlighting the active pocket (green color) **(c)** comparison of CLas PFP model with 3K2Q, *E.coli* PFK-1(PDB ID: 1PFK) (pink), PFK* from *Trypanosoma brucei* (PDB ID: 2HIG) (light blue), and the active pocket highlighted using green color in cartoon representation. The RFPEFR motif of 1PFK is highlighted in maroon, and the distinct loop regions of *C*Las PFP and 2HIG are shown in grey and light orange, respectively. (d) Predicted *C*Las PFP model in cartoon representation (blue), with the catalytic domain and the active-site residues of the PPi- and Mg2+-bound region shown on the right; hydrogen bonds in brown lines. The PFP+Mg^2+^ complex with F6P (**e**) and S7P (**f**) shows an enlarged view of interactions formed after a simulation run of 300 ns. The hydrogen bond interactions are shown as maroon lines.

Structural alignment of *C*Las PFP model with 3K2Q showed conservation of the catalytic domain with minimal deviation (**Figure 4b**). RMSD values for *C*Las PFP superimposed on 2HIG and 1PFK were 1.99 Å and 1.65 Å, respectively. A PDB similarity search ranked E. coli PFK-1 (1PFK) highest at 28% identity, followed by 3K2Q (27.3%) and 2HIG (25.4%). Superimposition with *E. coli* PFK-1 showed the ADP-binding motif RFPEFR (72-77), absent in other PFK variants. The RFPEFR motif of 1PFK structure adopts helical structure but forms a loop-and-sheet segment in the CLas PFP active site (**Figure 4c**). The 2HIG structure showed two distinct active-site loops: one containing the GGDG motif and the 332-348 loop, a conformation distinct from that of other bacterial PFKs. *C*Las PFP also has a unique 282-299 loop with a β-turn at DSFG (288-291), which is absent in 1PFK and missing from the 3K2Q crystal structure, probably due to weak density (**Figure 4c**). Thus, the results indicate that PFK variants use different secondary elements to accommodate their substrates/co-substrates.

The AlphaFold3 model of *C*Las PFP comprises α/β containing fold to accommodate the substrate, inorganic phosphate (PPi), and Mg^2+^. The secondary structure composition predicted using YASARA software was 47.2% helix, 18.0% sheet, 10.0% turn, and 22.2% coil. The active site residues predicted for PFP protein using NCBI conserved domain search tool were G12, G13, Y45, R79, D122, D123, T124, T126, T127, T150, D152, D154, M195, G196, R197, E267, Y324, and R327. The model (PFP+Mg^2+^) was docked with PPi and subjected to energy minimization using YASARA minimization server^[28]^. The energy minimized model of PFP+Mg^2+^ showed PPi occupying the catalytic pocket, forming interactions with G12, G13, Y45, S78, R79, V80, K81, D122, D123, T124, and K149, as well as polar contacts with R79, V80, K81, and T124. The Mg^2+^ ion is coordinated with D122, D152, and D154 (**Figure 4d**).

### 2.4 Molecular docking and simulation

Molecular docking of PFP+Mg^2+^ with the substrates F6P and S7P was conducted to assess the binding poses. Molecular docking revealed that F6P and S7P formed complexes with CLas PFP+Mg^2+,^ with binding energies of-5.7 and-5.9 kcal/mol, respectively. Molecular dynamics of these complexes and apo protein were carried out using GROMACS 2022.2 software [29]. Further, each complex bound to substrates after a 300ns simulation was analyzed. The active site residues involved in the binding of S7P with PFP+Mg^2+^ include G12, G13, R79, D122, D123, T124, T150, D152, R197 and R327. The substrate S7P formed polar contacts with the following residues: G13, I14, D122, T124, K149, and PPi in the binding pocket, thereby strengthening the interaction (**Figure 4f**). F6P formed interactions with the residues G12, G13, D122, T150, D152, and R327, along with PPi, in the binding pocket of PFP+Mg^2+^. Also, F6P forms polar contact with the residues G13, K149, and D152 (**Figure 4e).** Furthermore, the stable binding of substrates to the PFP+Mg^2+^ model was assessed. The Root Mean Square Deviation (RMSD) plotted with respect to backbone atoms depicted the average values of 0.32 nm, 0.19 nm, 0.24 nm, and 0.2 nm obtained for apo, PFP+Mg^2+^, F6P, and S7P, respectively (**Fig. S1a)**. The RMSD plotted with respect to ligand atoms showed the stable binding of the substrates F6P and S7P with an average value of 0.34 nm and 0.29 nm, respectively (**Fig. S1b)**. The average values of Solvent Accessible Surface Area (SASA) obtained for apo, PFP+Mg^2+^, F6P, and S7P were 199.1 nm^2^, 199.5 nm^2^, 202.1 nm^2^, and 196.4 nm^2^, respectively, implying the stable formation of the complexes (**Fig. S1c).** The average Root Mean Square Fluctuation (RMSF) values of 0.17 nm, 0.15 nm, 0.16, and 0.14 nm were attained for apo, PFP+Mg^2+^, F6P, and S7P, respectively (**Fig. S1d)**. The RMSF plot shows that the ligands formed a stable complex with the protein, with less fluctuation in the loop regions and active pocket residues. The average Radius of gyration (Rg) values obtained for apo, PFP+Mg^2+^, F6P, and S7P were 2.45 nm, 2.42 nm, 2.41 nm, and 2.37 nm, respectively. Thus, the binding of the substrate didn’t impact the packing of the protein (**Fig. S1e)**. The average number of hydrogen bonds formed by F6P and S7P with PFP was 2.9 and 4.5, respectively (**Fig. S1f)**.

The effective free energy or enthalpy of binding (ΔG_bind_) calculation using MMPBSA for the complexes PFP+Mg^2+^-F6P and PFP+Mg^2+^-S7P was-14.63 kcal/mol and-38.1 kcal/mol, respectively (**Table S3**). The per-residue decomposition analysis predicted the residues involved in the stable interaction formed by the ligands with PFP+Mg^2+^ (**Fig. S2**). The substrate F6P formed interaction with the residues A11, G12, G13, I14, K149, T150, I151, M195 and R327. In which R327 showed the lowest energy contribution of-1.68 kcal/mol, followed by I151, I14, and K149 with average energy values of-1.42 kcal/mol,-1.3 kcal/mol, and-1.23 kcal/mol, respectively (**Fig. S2a**). Compared with F6P, S7P formed more number interactions by A11, G12, G13, I14, R79, G120, G121, T124, K149, T150, I51, R327, Mg^2+^ and PPi. The average value of energy contributions shown by the residues K149> PPi> R79> I151> I14> T150> R327> G120 were-2.7 kcal/mol,-2.35 kcal/mol,-1.29 kcal/mol,-1.2 kcal/mol,-1.19 kcal/mol,-1.18 kcal/mol,-1.17 kcal/mol, and-1.03 kcal/mol, respectively (**Fig. S2b**).

### 2.5 Heterologous expression and purification of *C*Las PFP

The *C*Las PFP protein was heterologously expressed in *E.coli* Rosetta cells, and the overexpressed recombinant protein was purified using a pre-equilibrated Ni-NTA affinity column. The protein was eluted with 350 mM imidazole-containing elution buffer, dialyzed, and concentrated. The purity of the protein was assessed on 12% SDS-PAGE, and the purified protein showed a single band corresponding to a molecular mass of ∼44 kDa (**Fig. S3a**). The protein was further analyzed by size-exclusion chromatography (SEC), which showed a single peak corresponding to a molecular mass of ∼185.3 kDa, approximately four times the monomer mass of 44 kDa, indicating a homotetrameric assembly of the protein (**Fig. S3b**). The purified protein was pooled and further used for biochemical analysis.

### 2.6 Evaluation of the enzyme activity of *C*Las PFP

The enzyme activity of *C*Las PFP in phosphorylating F6P with different metal ions and phosphoryl donors was assessed. The enzyme assay showed that the protein has the highest relative activity for Mg^2+,^ followed by K^+^ (**Figure 5a**), and exhibits higher activity towards PPi than ATP. For further assay, Mg^2+^ was used as the metal cofactor and PPi as the phosphoryl donor. The *C*Las PFP was assessed for its enzyme activity on the substrates F6P and S7P to validate its dual functional role in the central carbon metabolism of *C*Las. The enzyme activity was examined at substrate concentrations ranging from 2.5 μM to 300 μM. The enzyme exhibited comparable substrate affinity towards S7P and F6P, with K_m_ values of 58 ± 3 μM and 67 ± 5 μM, respectively. The S7P and F6P showed catalytic efficiency with k_cat_/K_m_ values of 3.61 × 10^2^ M^-1^s^-1^ and 1.83 × 10^2^ M^-1^s^-1^, respectively (**Table 1)**. The results indicate that the enzyme showed affinity for both substrates, which fitted the Michaelis-Menten model (**Figures 5b and 5c**).

**Figure 5.**
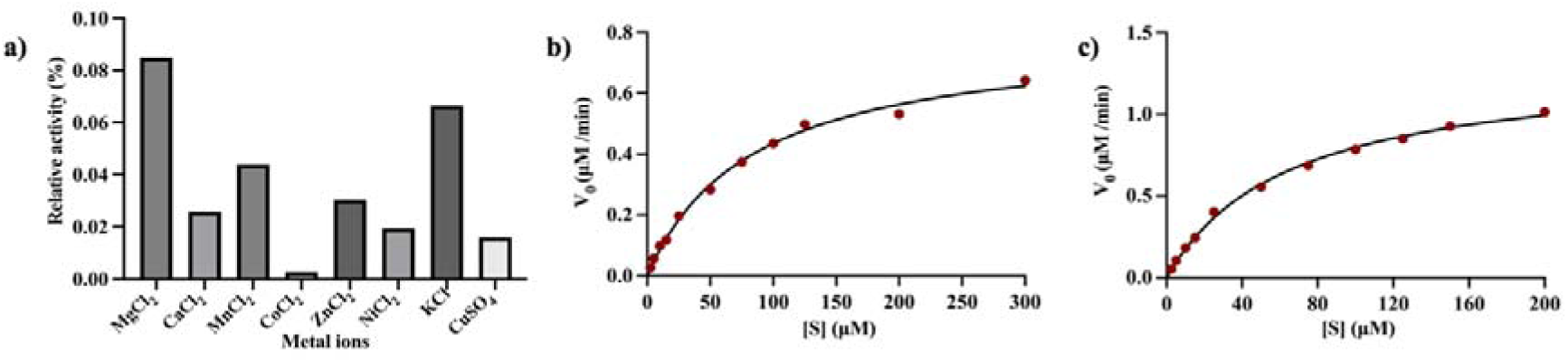
The enzyme kinetics of *C*Las PFP studied with the detection of release of inorganic phosphate from PPi on the phosphorylation of substrates using malachite green reagent. Kinetics: **a)** relative activity with different metal ions; Michaelis-Menten fits for the substrates **b)** F6P and **c)** S7P.

**Table 1.**
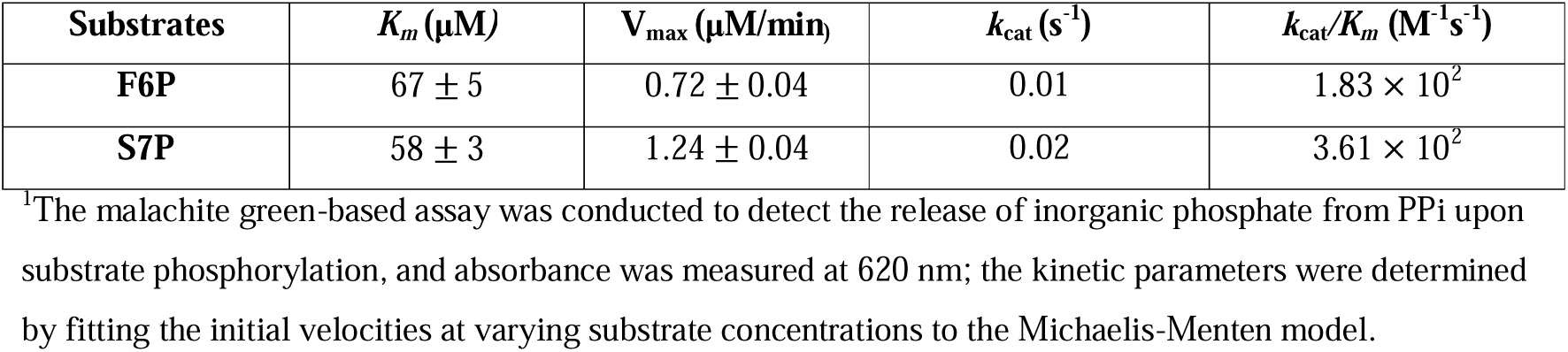
The kinetic parameters were calculated for 1 μM *C*Las PFP catalyzing the substrates in the presence of 2 mM MgCl_2_ and 2 mM PPi.

| **Substrates** | ***K<sub>m</sub>*** **(μM)** | **V<sub>max</sub> (μM/min<sub>)</sub>** | ***k*<sub>cat</sub> (s<sup>-1</sup>)** | ***k*<sub>cat</sub>/*K<sub>m</sub>*** **(M<sup>-1</sup>s<sup>-1</sup>)** |
|---|---|---|---|---|
| **F6P** | 67 ± 5 | 0.72 ± 0.04 | 0.01 | 1.83 × 10<sup>2</sup> |
| **S7P** | 58 ± 3 | 1.24 ± 0.04 | 0.02 | 3.61 × 10<sup>2</sup> |

<sup>1</sup>The malachite green-based assay was conducted to detect the release of inorganic phosphate from PPi upon substrate phosphorylation, and absorbance was measured at 620 nm; the kinetic parameters were determined by fitting the initial velocities at varying substrate concentrations to the Michaelis-Menten model.

### 2.7 Biophysical characterization of *C*Las PFP

CLas PFP was assessed at varying temperatures and pH by CD spectroscopy over the far-UV range 190–260 nm. The CD spectra showed pronounced negative bands at 208 nm and 222 nm, indicating predominant alpha-helical content. The thermal stability of the protein was investigated across the temperature range of 20 °C to 90 °C, showing the loss of alpha helices from 50 °C onward, with a prominent decrease observed by 60 °C (**Figure 6a**). From 20 °C to 60 °C, alpha-helical content decreased from 48.9% to 3.1%, with a concomitant increase in beta-sheet content from 11.5% to 38.1%. The helical content predicted for the AlphaFold3 model using YASARA was 47.2%. The observed melting temperature (Tm) of the protein was 51.1 ± 1.02 °C (**Figure 6c**). To assess conformational changes in CLas PFP upon incubation with each substrate, F6P and S7P showed only small changes relative to the apo protein. The observed Tm of the protein in the presence of F6P and S7P was 51.2 ± 2.3 °C and 51.8 ± 1.4 °C, respectively (**Fig. S4**).

**Figure 6.**
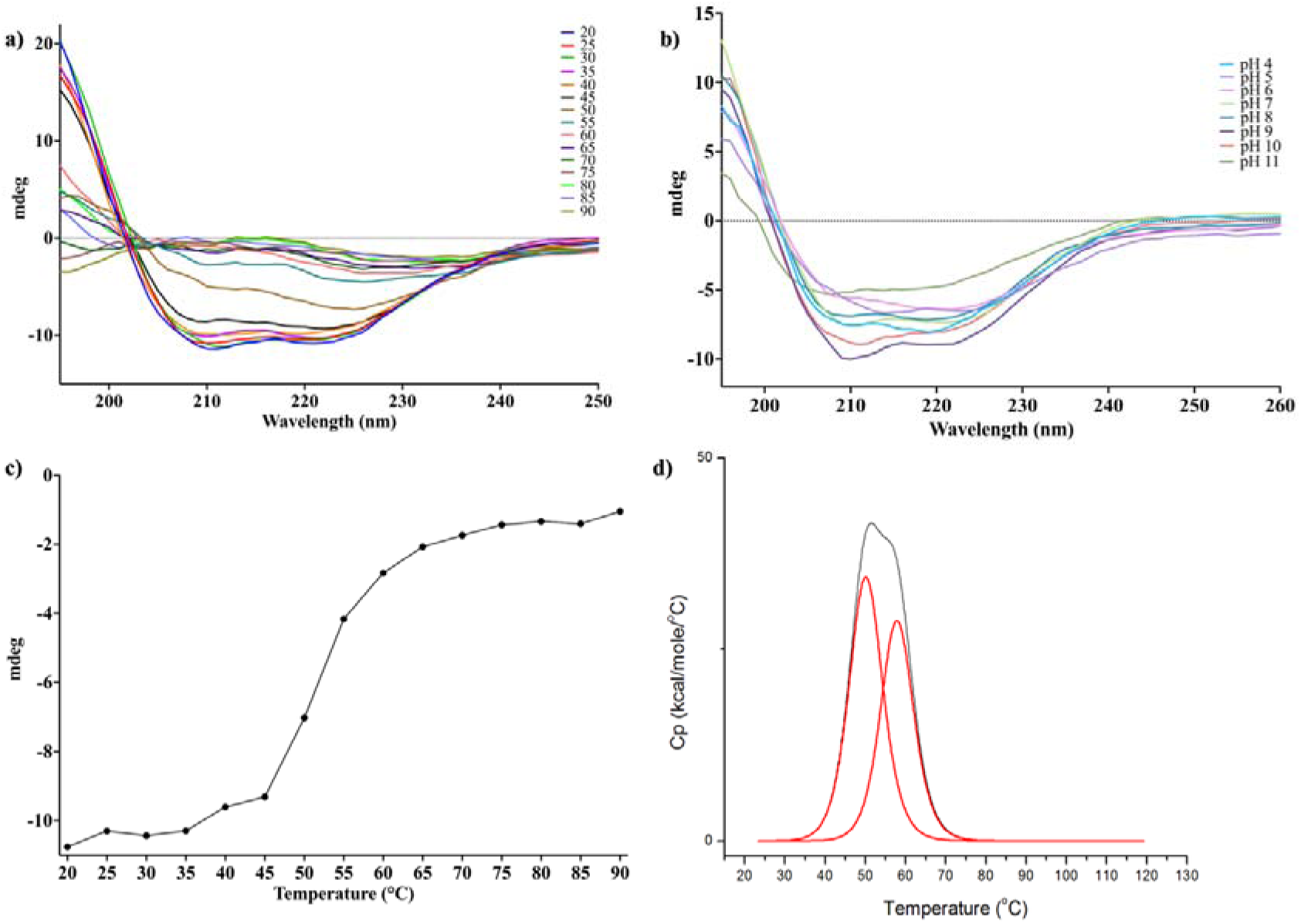
The biophysical characterization of *C*Las PFP protein. CD spectral analysis showing conformational transitions at a) different temperatures and b) different pH levels. c) The CD spectral signal (mdeg) at 222 nm tracing thermal unfolding. d) The thermal stability assessed using DSC.

Evaluating conformational changes across pH values from 4.0 to 11.0 revealed changes in secondary-structure conformations. At pH 4.0, the helical content was 42.9%, and the beta-sheet content was 13.6%. But at pH 5.0, the helical content decreased to 32.3%, and the beta-sheet content increased to 18.1%. Also, a gradual increase in helical content was observed as the pH increased from 6.0 to 9.0, from 38.3% to 45.9%, while beta-sheet content decreased from 16.9% to 10.9%. In the range of pH 9.0 to 11.0, protein lost alpha helical structure from 45.9% to 27.0% and increased beta sheet structure by 9%, i.e., 10.9% to 19.9% (**Figure 6b**). DSC showed two unfolding transitions, Tm_1_ 50.1 °C and Tm_2_ 57.9 °C (**Figure 6d**), suggesting that dissociation involves multistate unfolding or the formation of intermediates.

## 3. Discussion

In the course of evolution, host-dependent pathogens adopt a survival strategy that associates them with distinct host environments, leading to genome streamlining and functional adaptation to the host. Overdependence of *C*Las on the host has led to the loss of essential genes and disruption of metabolic pathways, including central carbon metabolism. In *C*Las, the absence of the genes encoding *pgi* and the PTS interrupts the glycolytic pathway from the synthesis of F6P. Also, the absence of transaldolase has blocked the non-oxidative branch of the PPP, preventing the interconversion of S7P into carbon intermediates that connect to the glycolytic pathway^[16,17]^. In the present study, we identified PFP belonging to the phosphofructokinase superfamily, which catalyzes dual reactions connecting the glycolytic and PPP pathways **(Figure 1)**. While there are reports showing the multifunctional role of PFP in other organisms^[30,31]^, our biochemical characterization of recombinant PFP validates this dual catalytic role in *C*Las.

The most canonical type of PFK is PFK-1, which converts F6P to FBP, whereas PFP catalyzes the equivalent function using PPi and thereby conserves ATP^[30]^. Consistent with this, CLas PFP was more active with PPi than with ATP. Earlier work has reported horizontal gene transfer among PFKs and active-site mutations that shift phosphoryl donor preference^[32]^. The ATP/PPi binding site is defined by the signature motifs GG(D/N)(**D/G)** and (**K/G)**TIDXD, in which the two glycines conserved in PFK-1 are replaced by aspartate and lysine in PFP **(Figure 2).** Mutation of the corresponding residues from this motif to glycine (D175→G) and (K201→G) from *E. histolytica* PFP showed activity towards ATP on double mutation and lost activity with PPi^[33]^. This suggests that ATP-binding activity is encoded inherently in the gene^[33]^. Evolutionary analysis showed that other variant clades share the same LCA as PFP, suggesting that PFP may be ancestral (**Figure 3**). Previous studies proposing PFK* or PFK-1 as the ancestral protein remain in conflict^[32,34]^. However, unlike PFK-1, PFP catalyzes a reversible reaction, lacks an allosteric site of regulation, and plays a multifunctional role, suggesting that PFP could be ancestral^[30]^. The present data are insufficient to conclude that PFP is ancestral; however, our result infers the possibility.

The predicted model of *C*Las PFP adopts an α/β fold accommodating PPi and Mg^2+^ **(Figure 4d)**. The crystal structures of PFK proteins from different species have been shown to have two or three domains characterized as Rossmann-like fold, forming a three-layer sandwich (α-β-α)^[35,36]^. *C*Las PFP differs from PFK-1 and PFK* at the ATP/PPi binding site. The residues involved in the binding of the nucleotide moiety in the active pocket of *E. coli* PFK-1 were Y41, F76, R77, R82, and R111^[37]^. The adenine-binding motif RFPEFR (72–77) of PFK-1 is absent in *C*Las PFP. This RFPEFR motif, represented in a 3_10_ helix ^[35]^, is modulated into a loop segment in the PFP and PFK* protein structures (**Figure 4c).** G108 of 1PFK, key to ATP binding, is replaced by threonine (T127) in CLas PFP, altering pocket hydrophobicity^[36]^. The PFK* with motifs GGD**G** and **K**TIDXD was previously reported in kinetoplasts, *Leishmania donovani* and *T. brucei* belonging to the eukarya domain^[38]^. T. brucei PFK* has a unique N-terminal domain and a 329–348 loop adjacent to the active pocket, proposed to contribute to ATP binding^[39]^. Interestingly, CLas PFP structure showed a distinct 282-299 loop region, which is missing from other compared structures (**Figure 4c)**. But a similar loop (380–390) was reported in one of the subunits of PFP from *Borrelia* burgdorferi and may be part of substrate binding^[36]^.

CD spectroscopy was used to assess structure and conformational stability. The CD results complement the homology model, showing predominant contributions from alpha helices. Helical content was retained up to 45 °C, unfolding began at 50 °C, and secondary structure was lost by 60 °C (**Figure 6a**). The thermal denaturation studies conducted on PFK protein also showed that major structural transitions occur between 55 °C and 60 °C^[40]^. The Tm obtained from the CD spectral analysis, 51.1 ± 1.02 °C, is in agreement with the first DSC transition (50.1 °C) (**Figure 6c & 6d**). The thermal stability study of PFP with the substrates F6P and S7P showed a slight increase in Tm, and the helical content began to decrease after 50 °C (**Fig. S4**). The structural analysis of substrate binding showed minimal changes in the loop region and active site residues, suggesting that only minor structural changes are induced in the PFP structure upon substrate binding (**Fig. S1d**).

Across the pH range 4.0 to 11.0, helical content increased as pH increased from 5.0 to 9.0. A study on the PFK-1 protein assessing the transition at varying pH has shown the shift in the dimer-tetramer equilibrium as pH increased from 6.0 to 8.0. Additionally, phosphate has been shown to influence the increase in helical content with increasing pH ^[40]^. Similar to these results, *C*Las PFP may shift to the tetrameric/oligomeric state as pH increases, although the oligomeric state was not assessed here. *C*Las is associated with two hosts: citrus phloem, which was found to be slightly acidic, with a pH of ∼6.0 [41], and psyllid hemolymph, which showed an increase in pH to ∼8.0 after CLas infection[42]. This defines the physiological pH range in which the protein can remain stable, between 6.0 and 8.0. The optimal pH range for PFP homologs falls between 6.0 and 8.0[43]. The CD spectra also showed the loss of helical structure observed at pH 11.0, indicating a loss of structural integrity, and pH 11 is far from the physiological range (**Figure 6b)**.

To evaluate the activity of recombinant *C*Las PFP towards different co-substrates, a malachite green-based assay was performed with F6P as the substrate. CLas PFP showed higher activity with PPi than ATP, confirming PPi as the preferred phosphoryl donor, and was most active with Mg^2+^, followed by K^+^ (**Figure 5a**). The reaction mechanism reported for PFK involves the nucleophilic attack of 1-OH group of F6P on the terminal phosphoryl group of ATP/PPi, thus facilitating substrate phosphorylation^[36]^. During cleavage of the phosphoanhydride bond, the terminal phosphate adopts a pentacoordinate state, and Mg^2+^ plays an essential role in stabilizing this bond^[35]^. As proposed earlier, monovalent ions, such as K^+^ or NH4+, can also substitute for Mg^2+[35]^.

The functional annotation of *C*Las genome revealed that PFP protein is involved in the catalysis of two reactions, F6P to FBP (glycolysis) and S7P to SBP (PPP) (**Figure 1**). Steady-state kinetics of the recombinant enzyme confirmed conversion of both F6P and S7P, with K_m_ values of 67 ± 5 μM and 58 ± 3 μM, respectively (**Table 1**). Previous studies investigating PFP from *Entamoeba histolytica* showed substrate affinity with K_m_ values of 64 μM and 38 μM towards S7P and F6P, respectively^[20]^. On comparing, *E. histolytica* showed more substrate affinity towards F6P, and CLas PFP prefers S7P than F6P. Additionally, *Clostridia spp.* that lack transaldolase have been shown to use PFP to catalyze both S7P and F6P with high affinity, unlike *E. coli* PFK-1^[31]^. Thus, the biochemical analysis confirmed the presence of a functional PFP protein with dual specificity that catalyzes the forward reaction. Bacterial PFP are non-allosteric and are primarily regulated by PPi/Pi ratios and relative substrate concentrations [44]. It has been reported that PFP is reversible, catalyzing the conversion of FBP to F6P and SBP to S7P, with the concomitant conversion of Pi to PPi. PFP from *Methylomicrobium alcaliphilum* has been shown to catalyze the reaction for both F6P (K_m_ = 0.328 mM) and S7P (specific activity of 22 U mg^-1^), supporting the bidirectionality of PFP [45].

*In silico* structural analysis showed stable binding of PFP to both substrates. The interaction study results infer catalytic residues involved in the binding of F6P and S7P. The interaction study showed that K149 forms a hydrogen bond with F6P and S7P (**Figure 4e, 4f**). The PFP protein of Borrelia burgdorferi (PDB ID:1KZH) proposed that K203 (corresponding to K149 in CLas PFP) plays an important role in the coordination of PPi and stabilization of the transition state in PFP proteins [36] It forms a stable interaction with an energy of-1.23 kcal/mol and-2.7 kcal/mol, respectively (**Fig. S2**). Also, in 1PFK, this residue is substituted with glycine at position 124, and this reaction mechanism involving lysine can be unique to PFP proteins. These motif differences provide a plausible structural basis for the distinct phosphoryl-donor preferences of the PFK groups (**Figure 2**). In PFP, the aspartate of the GG(D/N)D motif (D123 in CLas PFP; D177 in B. burgdorferi PFP) is positioned to exclude the α-phosphate of ATP, while the lysine of the KTIDXD motif (K149) coordinates PPi [36]. In ATP-dependent PFK-1, both positions are occupied by glycine (GGDG and GTIDXD), thereby creating space for the nucleotide and removing the PPi-binding lysine, consistent with ATP specificity [35,36]. PFK* combines the ATP-type GGDG motif with the PPi-type KTIDXD motif, and its ATP dependence suggests that loss of the motif aspartate alone is sufficient to permit ATP binding [39]. Likewise, the GGDN motif of PFK# and the substitutions at R197 (Q/E/H) may further modulate donor or sugar-phosphate binding. However, these relationships are inferred from sequence and structural comparisons, and kinetic characterization of the variants is needed to confirm them.

One of the multifunctional roles of the PFP protein is its involvement in the SBPP pathway, which converts S7P to SBP and can bypass the loss of transaldolase by working together with FBPA. Variants of this pathway have been reported in the human gut bacterium *Prevotella copri*^[21,22]^, *Entamoeba histolytica^[20]^,* and *cellulolytic clostridia^[31]^.* Interestingly, kinetic studies confirmed that *C*Las PFP can also catalyze the reaction to form SBP from S7P. So, the question arises: Is *C*Las capable of bypassing the loss of transaldolase? As mentioned in previous research, the SBP metabolite could be overlooked, or the misannotation of FBPA as transaldolase in organisms could explain the limited identification of the SBP pathway across species^[31]^. Additionally, a multiple-knockout study involving *E. coli* transaldolase mutants revealed that bacteria depend on PFK-1 and FBPA to complete PPP^[46]^. This outcome suggests that the SBP pathway could be a compensatory route or survival strategy adopted by species with disrupted metabolic function. Hence, it is possible that *C*Las, which lacks transaldolase, uses SBPP as an alternative route, or that PFP may serve an alternative function in *C*Las metabolism. *C*Las imports ATP directly from the host using ATP translocase. If SBPP pathway is active in *C*Las, it raises the possibility that the pathogen reroutes interrupted glycolysis to produce ATP via substrate-level phosphorylation under certain conditions, such as limited availability of host ATP. The present study demonstrates the catalytic efficiency of recombinant *C*Las PFP; however, further studies involving *C*Las FBPA and *in planta* experiments are required to verify the presence of SBPP in *C*Las.

## 4. Methodology

### 4.1 Genome analysis and Functional annotation

The genome and coding sequence (CDS) of *C*Las strain gxpsy (unculturable) (NC_020549.1) and Lcr strain BT-1 (culturable) (NC_019907.1) were fetched from Genbank, NCBI^[47]^. The genome annotation of Lcr BT-1 and *C*Las was carried out using the RAST annotation server^[48]^, and the resulting genome sequence data were deployed to build a metabolic model using the Kbase platform^[49]^. The two metabolic models were compared for reactions in central carbon metabolism, and the results were manually curated using protein-protein BLAST 2.14.0^[50]^ with an e-value cutoff of 1e-05.

Carbon metabolism genes missing or unique (non-canonical reactions) in *C*Las were then compared with homologs to assess how pathways differ across species. Genomes and coding sequences (CDSs) of 35 homologs from the phyla proteobacteria, bacteroidota, firmicutes, and evosea were retrieved from NCBI, and BLASTP search against a database derived from their CDS sequences were conducted for three major carbon metabolism-related genes.

### 4.2 Sequence analysis

Functional annotation identified a key reaction catalyzed by PFP, whose nucleotide and protein sequences (WP_015452326.1) were retrieved from NCBI. The active site residues of PFP were predicted using NCBI conserved domain search^[23]^. Sequences of phosphofructokinase (PFK) variants were retrieved from OrthoDB, supplemented by BLASTP against ClusteredNR; 52 sequences were included, spanning the bacterial phyla proteobacteria, bacteroidota, actinomycetota, and firmicutes and the eukaryotic phyla evosea, euglenozoa, and apicomplexa. The multiple sequence alignment (MSA) was performed for the selected sequences using the MAFFT program^[51]^ with the FFT-NS-i iterative refinement method and BLOSUM62 for amino acid sequences and PAM1 for nucleotide sequences. A maximum-likelihood tree was constructed from the aligned PFP and homolog sequences in MEGA 11^[24]^, with 1000 bootstrap replicates. The tree was visualized using iTOL v7^[52]^.

### 4.3 Structure prediction

The three-dimensional structure of *C*Las PFP was predicted using AlphaFold3^[25]^, ESMFold^[26]^, and RoseTTAFold^[27]^. The PFP protein bound to Mg^2+^ ion was modeled using AlphaFold3. Homologous structures were obtained by BLASTP against the PDB using *C*Las PFP as the query, and predicted structures were compared with *Marinobacter nauticus* PFP (PDB ID: 3K2Q) to select the best model by lowest RMSD. The modeled proteins were further validated by checking stereochemistry using PROCHECK in the SAVES server v6.0^[53]^. The validated model was compared with other homologous structures and docked with the co-substrate pyrophosphate (PPi) using AutoDock Vina^[54]^ and refined using energy minimization in the YASARA minimization server^[28]^. The final model was visualized and analyzed in UCSF ChimeraX software^[55]^.

### 4.4 Molecular docking and structure refinement

The two-dimensional structures of substrates fructose 6-phosphate (F6P) and sedoheptulose 7-phosphate (S7P) were retrieved from PubChem [56]. Both protein and ligand structures are pre-processed, and rigid docking was carried out using Autodock Vina in AMDock 1.5.2 graphical tool [54]. Based on the binding affinity, the best docked poses were selected, and interaction analysis was performed in PyMOL. Molecular dynamics of the apo protein and protein-ligand complex [57] were carried out using GROMACS software. The best ligand poses obtained from docking were assigned with H atoms using Avogadro software [58]. The ligand topology and parameterization were performed using the CHARMM General Force Field (CGenFF) webserver [59]. In contrast, the CHARMM forcefield and TIP3P water module of GROMACS were utilized to generate protein topology. The initial complex of the system was generated by combining protein (PFP), cofactor (PPi and Mg^2+^), and respective substrate (F6P/S7P) structures and corresponding topology were appended. A unit cell of a dodecahedron shape with a 2.0 nm padding centering protein-ligand complex was generated, and an SPC module was used to solvate the system. Further, the neutralization of the system was done by adding Na^+^/Cl^-^ ions and utilizing the steepest descent method to perform energy minimization for 100ps, converging to a maximum force of 1000 kJ/mol/nm. The NVT and NPT equilibration steps were carried out for 300 K and 1 bar pressure, respectively. Finally, the system was subjected to a 300 ns simulation run. The parameters used to elucidate structural stability include RMSD with respect to backbone atoms and ligand atoms, RMSF, Hydrogen bond formation, SASA, and Rg. To calculate the effective free energy of binding of PFP+Mg^2+^-substrate complexes, Molecular Mechanic/Poisson-Boltzmann Surface Area (MMPBSA) method was used by gmx_MMPBSA v1.6.4 and AmberTools-20 [60]. The last 1000 frames of the simulation run were taken to calculate the binding energy for a multi-component system with PFP+Mg^2+^ as the receptor and the substrate molecule as the ligand. The per-residue decomposition analysis was performed to predict the key residues involved in the binding.

### 4.5 Heterologous expression and purification of *C*Las PFP

The recombinant plasmid of *C*Las PFP gene in vector pET-28c was synthesized by Gene Bio Solutions (India) and transformed into *E. coli* Rosetta cells. Gene expression was induced with 0.4 mM isopropyl β-d-thiogalactopyranoside (IPTG) at 16 °C for 16 h. Cells from a 1 L culture were resuspended in lysis buffer (50 mM Tris, 300 mM NaCl, 10% glycerol, pH 7.4) with lysozyme (0.1 mg/ml), sonicated for 15 min, and centrifuged at 10,000 rpm for 45 min. The supernatant was loaded onto a pre-equilibrated Ni²⁺-NTA column, and the protein was eluted with 350 mM imidazole buffer. The eluted protein was dialyzed using dialysis buffer (50 mM Tris, 150 mM NaCl, 10% glycerol, pH 7.4) at 4 °C for 10 h and concentrated using an Amicon Ultra concentrator (3 kDa cutoff). The protein concentration was determined by BCA assay (OD₅₆₂). The purity of the protein was assessed at each step on a 12% SDS-PAGE gel. Further, the purified protein was loaded onto a HiLoad Superdex 200 pg column at a flow rate of 0.5 mL/min to assess the purity and molecular mass and oligomeric state of protein. The SEC column was pre-calibrated with standard molecular weight markers including Thyroglobulin (660 kDa), Ribonuclease (440kDa), aldolase (158 kDa), Conalbumin (75kDa), Ovalbumin (44 kDa), Ribonuclease A (13.7 kDa) and Aprotinin (6.5 kDa).

### 4.6 Biochemical characterization of PFP

The specificity of *C*Las PFP towards different metal ions and phosphoryl donors was studied by detecting the release of inorganic phosphate from PPi/ATP during the phosphorylation of F6P (Cat. No./ID: F3627) (the natural substrate) using Malachite green reagent (0.03% (w/v) malachite green, 0.2% (w/v) ammonium molybdate and 0.05% (v/v) Triton X-100 in 0.7 N HCl) [61]. Each reaction mixture, consisting of 1 μM CLas PFP protein, 300 μM F6P, 2 mM PPi, and 2 mM of the different metal ions, including MgCl_2_, CaCl_2_, MnCl_2_, CoCl_2_, ZnCl_2_, NiCl_2_, CuSO_4,_ and KCl, was incubated for 10 minutes at 20 °C, with a blank control lacking protein for each metal ion. Malachite green was then added to all wells, and the wells were incubated at 20 °C for 20 minutes. Absorbance was measured at 620 nm to evaluate relative activity. For phosphoryl donor specificity, 2 mM PPi or ATP (Cat. No./ID: A2383) was added to parallel reaction mixtures and relative activity assessed. Phosphorylation of F6P and S7P (Cat. No./ID: 78832) was then examined in 100 μL reactions containing 1 μM *C*Las PFP, varying substrate concentrations, 2 mM MgCl_2_, and 2 mM PPi. The enzyme assay was conducted in triplicate, and the corresponding kinetic parameters were calculated by fitting the initial velocities to the Michaelis-Menten model.

### 4.7 Biophysical characterization of *C*Las PFP

Structural stability was assessed using a JASCO-1500 circular dichroism spectrophotometer (JASCO, Tokyo, Japan) fitted with a Peltier thermostat, recording spectra over 190–260 nm for 1 μM CLas PFP in a 1 mm quartz cuvette. The protein was incubated for 10 minutes under different pH (4.0 – 11.0) and temperature (20-90 °C) conditions, and conformational stability was evaluated. The conformational stability was evaluated in the presence of each substrate, F6P and S7P for 5 μM CLas PFP, under different temperatures (20-90 °C). Thermal stability was assessed by monitoring ellipticity at 222 nm against temperature, with three consecutive scans per condition. The smoothened CD data in millidegrees (mdeg) were converted to Mean Residual Ellipticity (MRE), and secondary structure content (%) was estimated using the K2D3 server^[62]^.

Thermal unfolding was measured by differential scanning calorimetry (DSC) on a MicroCal VP-DSC microcalorimeter (MicroCal, USA). Concentrated 50 μM *C*Las PFP and 25 mM Tris buffer were filtered, degassed, and loaded into the sample and reference cells, respectively. Experiments were conducted at 20–120 °C at 1 °C min^-1^. Scans were plotted and analyzed in Origin 7.0.

## Acknowledgment

SKP thanks Ministry of Education (MHRD), Government of India, for supporting with the fellowship. The authors are grateful to the computational facility provided by the Bioinformatics Center (BIC) and Ashok Soota Molecular Medicine Facility at IIT Roorkee.

## CRediT author statement

**Sandra Kathott Prakash**: Conceptualization, Data curation, Formal analysis, Methodology, Investigation, Validation, Visualization, Writing – original draft. **Etisha Arun & Khushhal Rathore:** Formal analysis, Methodology, Investigation, Validation, Visualization. **Mrugendra Gubyad**: Formal analysis, Methodology, Investigation. **Dilip Kumar Ghosh**: Formal analysis, Validation, Resources. **Jitin Singla & Ashwani Kumar Sharma**: Conceptualization, Validation, Supervision, Writing – Review and Editing.

## Declaration of generative AI and AI-assisted technologies in the manuscript preparation

During the preparation of this work, the author(s) used Claude Opus 4.8 for language editing and grammar checking. After using this tool/service, the author(s) reviewed and edited the content as needed and take(s) full responsibility for the content of the published article.

## Preprint

An earlier version of this manuscript was deposited as a preprint on bioRxiv ^[63]^.

