## Supplementary for "Identification and characterization of bifunctional pyrophosphate-fructose-6-phosphate 1-phosphotransferase from citrus greening disease-causing α-proteobacterium"

**Table S1.** The functional annotation results showing the number of annotated genes present in *C*Las gxpsy and *Lcr* BT-1 for each metabolic pathway. A lower number of genes, or their absence, is highlighted in bold.

| **Metabolism** | | **No. of genes in *C*Las gxpsy** | **No. of genes in *Lcr* BT-1** |
| --- | --- | --- | --- |
| **Carbohydrate metabolism** | Glycolysis | **12** | 13 |
|  | Pentose phosphate pathway | 8 | **7** |
|  | Pyruvate metabolism | 5 | 5 |
|  | Citric acid cycle | 15 | 15 |
|  | Methylglyoxal metabolism | **1** | 4 |
| **Energy metabolism** | Oxidative phosphorylation | **33** | 36 |
| **Amino acid metabolism** | Alanine, serine, and glycine | 6 | 6 |
|  | Arginine; urea cycle, polyamines | 9 | 9 |
|  | Aromatic amino acids and derivatives | **-** | 19 |
|  | Glutamine, glutamate, aspartate, asparagine; ammonia assimilation | 6 | 6 |
|  | Lysine, threonine, methionine, and cysteine | **16** | 20 |
|  | Histidine | **-** | 10 |
|  | Proline | **-** | 4 |
| **Lipid metabolism** | Fatty acid biosynthesis | 13 | 13 |
|  | Fatty acid degradation | **-** | 1 |
|  | Glycerolipid metabolism | **11** | 12 |
|  | Acyl-CoA hydrolysis | **1** | - |
| **Vitamin & cofactors, prosthetic groups, pigments** | Thiamin | **2** | 7 |
|  | Riboflavin | 5 | 5 |
|  | Pyridoxal 5-phosphate | 5 | 5 |
|  | Niacin | **5** | 6 |
|  | CoA biosynthesis | **5** | 7 |
|  | Biotin | 5 | 5 |
|  | Folate | **8** | 13 |
|  | Ubiquinone | 5 | 5 |
| **Nucleotide metabolism** | Purine | 16 | 16 |
|  | Pyrimidine | 14 | 14 |

**Table S2.** The presence (+) and absence (-) of central carbon metabolism-related enzymes in homologous species. PFK represents any phosphofructokinase isoforms belonging to phosphofructokinase type A (PFKA; IPR050929). PFP denotes the PPi-dependent phosphofructokinase, and pgi is phosphoglucoseisomerase.

| **Accession ID** | **Species** | **pgi** | **PFK** | **Transaldolase** | **PFP** |
| --- | --- | --- | --- | --- | --- |
| NC_015684.1 | *Afipia carboxidovorans OM5* | **+** | **-** | **+** | **-** |
| NZ_CP033022.1 | *Agrobacterium fabrum strain 1D132* | **+** | **+** | **+** | **+** |
| NC_009937.1 | *Azorhizobium caulinodans ORS 571* | **+** | **-** | **+** | **-** |
| NC_008783.1 | *Bartonella bacilliformis KC583* | **+** | **-** | **+** | **-** |
| NC_005956.1 | *Bartonella henselae str. Houston-1* | **+** | **-** | **+** | **-** |
| NC_005955.1 | *Bartonella quintana str. Toulouse* | **+** | **-** | **+** | **-** |
| NC_006932.1 | *Brucella abortus bv.1str.9-941* | **+** | **-** | **+** | **-** |
| NC_009667.1 | *Brucella anthropi ATCC 49188* | **+** | **-** | **+** | **-** |
| NC_003317.1 | *Brucella melitensis bv.1str.16M* | **+** | **-** | **+** | **-** |
| NZ_CP004021.1 | *Candidatus Liberibacter africanus strain PTSAPSY* | **-** | **+** | **-** | **+** |
| NC_022793.1 | *Candidatus Liberibacter americanus strain Sao Paulo* | **-** | **+** | **-** | **+** |
| **NC_020549.1** | ***Candidatus Liberibacter asiaticus strain gxpsy*** | **-** | **+** | **-** | **+** |
| NC_014774.1 | *Candidatus Liberibacter solanacearum strain CLso-ZC1* | **-** | **+** | **-** | **+** |
| CP120448.1 | *Candidatus Phytoplasma asteris* | **+** | **+** | **-** | **-** |
| NZ_CP120361.1 | *Chelativorans sp. AA-79* | **+** | **+** | **+** | **+** |
| AP023109.1 | *Entamoeba histolytica HM-1:IMSS* | **+** | **+** | **-** | **+** |
| NC_000913.3 | *Escherichia coli str.K-12substr.MG1655* | **+** | **+** | **+** | **-** |
| NZ_CM002917.1 | *Hoeflea phototrophica DFL-43* | **+** | **+** | **+** | **+** |
| **NC_019907.1** | ***Liberibacter crescens strain BT-1*** | **+** | **+** | **+** | **+** |
| NC_007626.1 | *Magnetospirillum magneticum AMB-1* | **+** | **+** | **+** | **+** |
| NC_002678.2 | *Mesorhizobium japonicum MAFF 303099* | **+** | **+** | **+** | **+** |
| NC_010505.1 | *Methylobacterium radiotolerans JCM 2831* | **+** | **-** | **+** | **-** |
| NC_011666.1 | *Methylocella silvestris BL2* | **+** | **-** | **+** | **-** |
| NC_010172.1 | *Methylorubrum extorquens PA1* | **+** | **-** | **+** | **-** |
| NC_007964.1 | *Nitrobacter hamburgensis X14* | **+** | **+** | **+** | **+** |
| NZ_CP085932.1 | *Prevotella copri DSM 18205strain FDAARGOS_1573* | **+** | **+** | **-** | **+** |
| NZ_CP021850.1 | *Pseudoclostridium thermosuccinogenes strain DSM 5807* | **+** | **+** | **-** | **+** |
| NC_021505.1 | *Pseudomonas putida NBRC 14164* | **+** | **-** | **+** | **-** |
| NC_008380.1 | *Rhizobium johnstonii 3841* | **+** | **+** | **+** | **+** |
| NC_014664.1 | *Rhodomicrobium vannielii ATCC* | **+** | **-** | **+** | **-** |
| NZ_CP116810.1 | *Rhodopseudomonas palustris CGA009* | **+** | **-** | **+** | **-** |
| NZ_CM011002.1 | *Roseibium alexandrii DFL-11* | **+** | **+** | **+** | **+** |
| NC_003047.1 | *Sinorhizobium meliloti 1021* | **+** | **+** | **+** | **+** |
| NZ_CP013197.1 | *Spiroplasma citri strain R8-A2* | **+** | **+** | **-** | **-** |
| NC_014217.1 | *Starkeya novella DSM 506* | **+** | **-** | **+** | **-** |

**Table S3.** The summary of effective free energy calculations for ligands, F6P and S7P, bound with PFP+Mg^2+^ using MMPBSA.

|  | **Interaction energy: ΔE_MM_ (kcal/mol)** | | **Solvation energy: ΔG_Solv_ (kcal/mol)** | | **Effective free energy: ΔG_bind_ (kcal/mol)** |
| --- | --- | --- | --- | --- | --- |
| **Ligands** | Van der Waal energy, ΔE_VdW_ | Electrostatic energy, ΔE_Ele/_ ΔE_Scf_ | Polar component, ΔG_GB/PB_ | Non-polar component, ΔG_non-polar_ | Enthalpy, ΔH, (kcal/mol) |
| S7P | -30.76 | -111.12 | 107.01 | -3.22 | -38.1 |
| F6P | -25.16 | -61.18 | 74.52 | -2.81 | -14.63 |

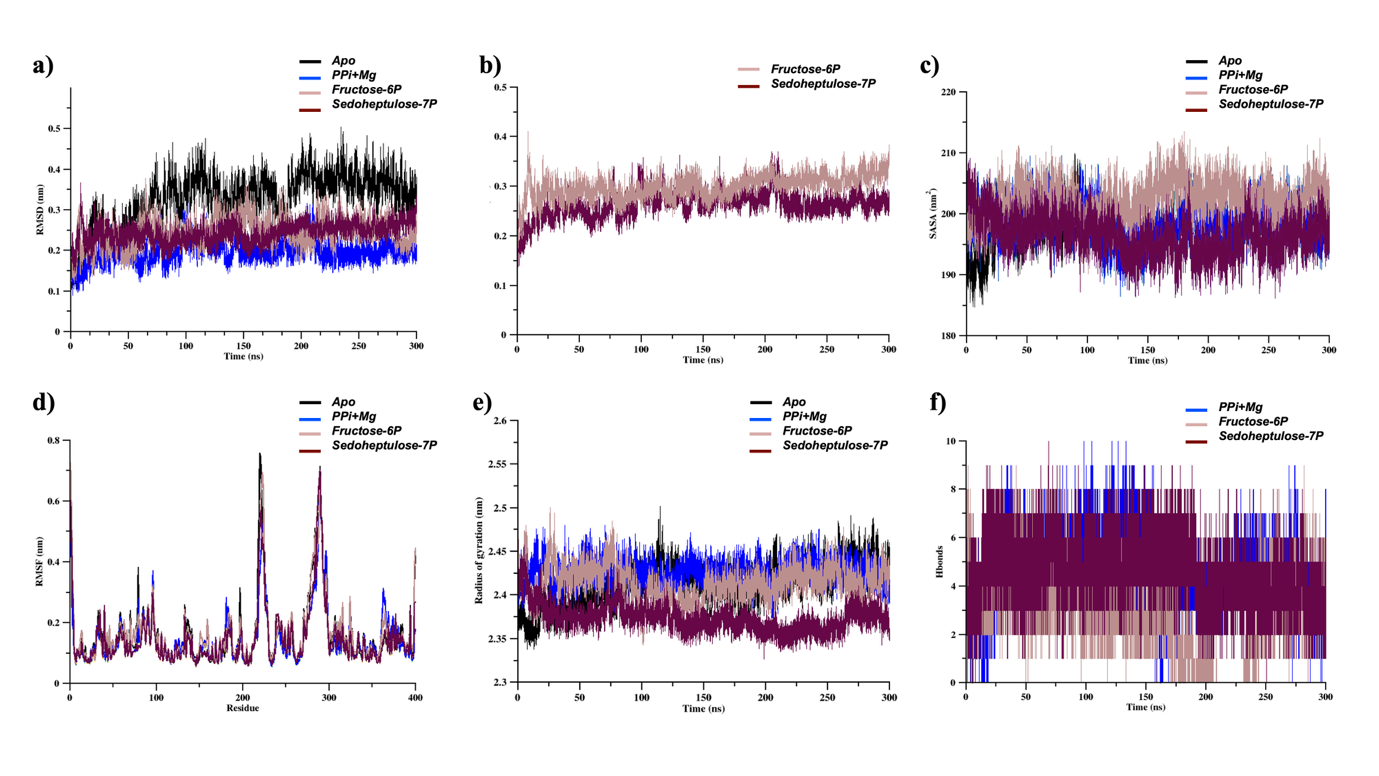

**Figure S1.** The structural stability analysis result after the simulation run for 300 ns. The plots of (**a**) RMSD-Backbone atoms, (**b**) RMSD-ligand atoms, (**c**) SASA, (**d**) RMSF, (**e**) Rg, and (**f**) number of H bond interactions represent the stability of PFP and complexes throughout the trajectories.

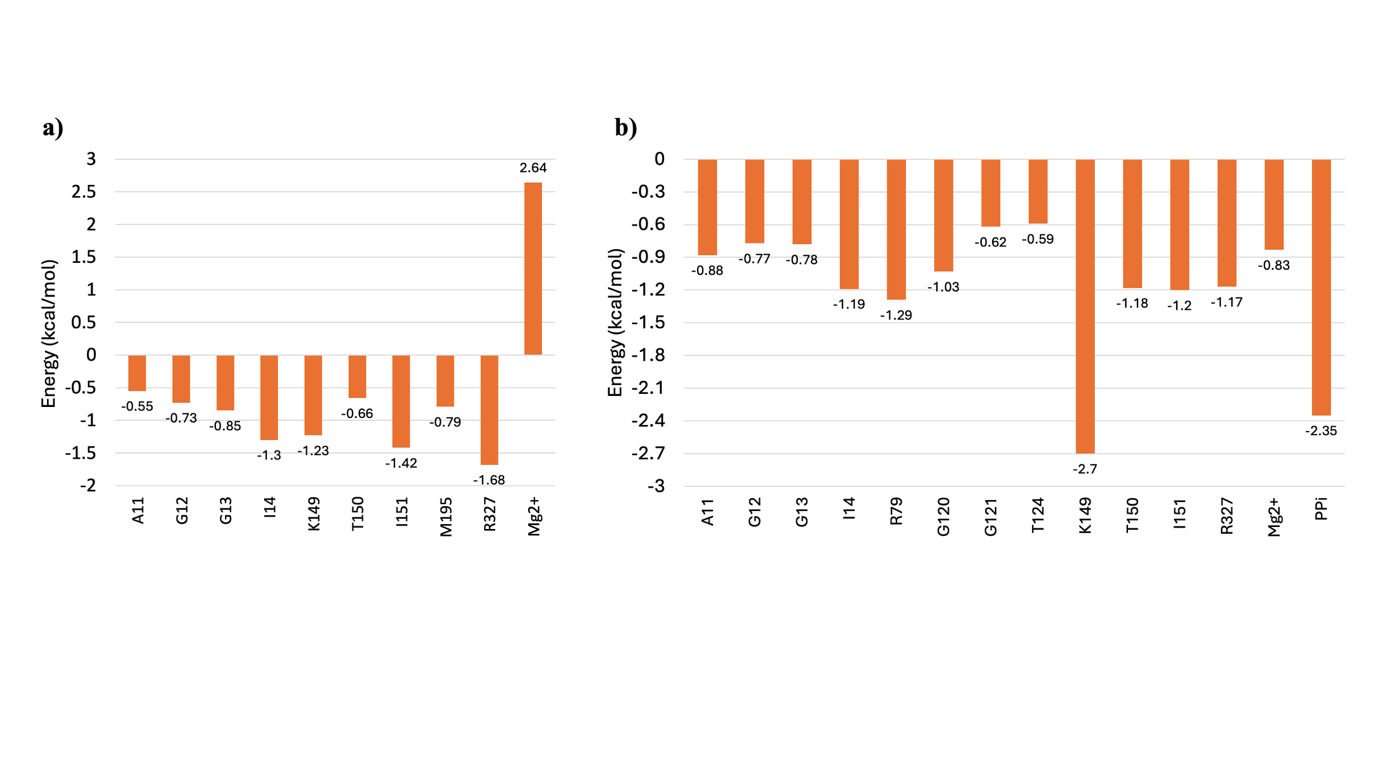

**Figure S2.** Per-residue decomposition analysis depicting the residues forming stable interaction with ligands a) F6P and b) S7P and corresponding energy contribution in kcal/mol.

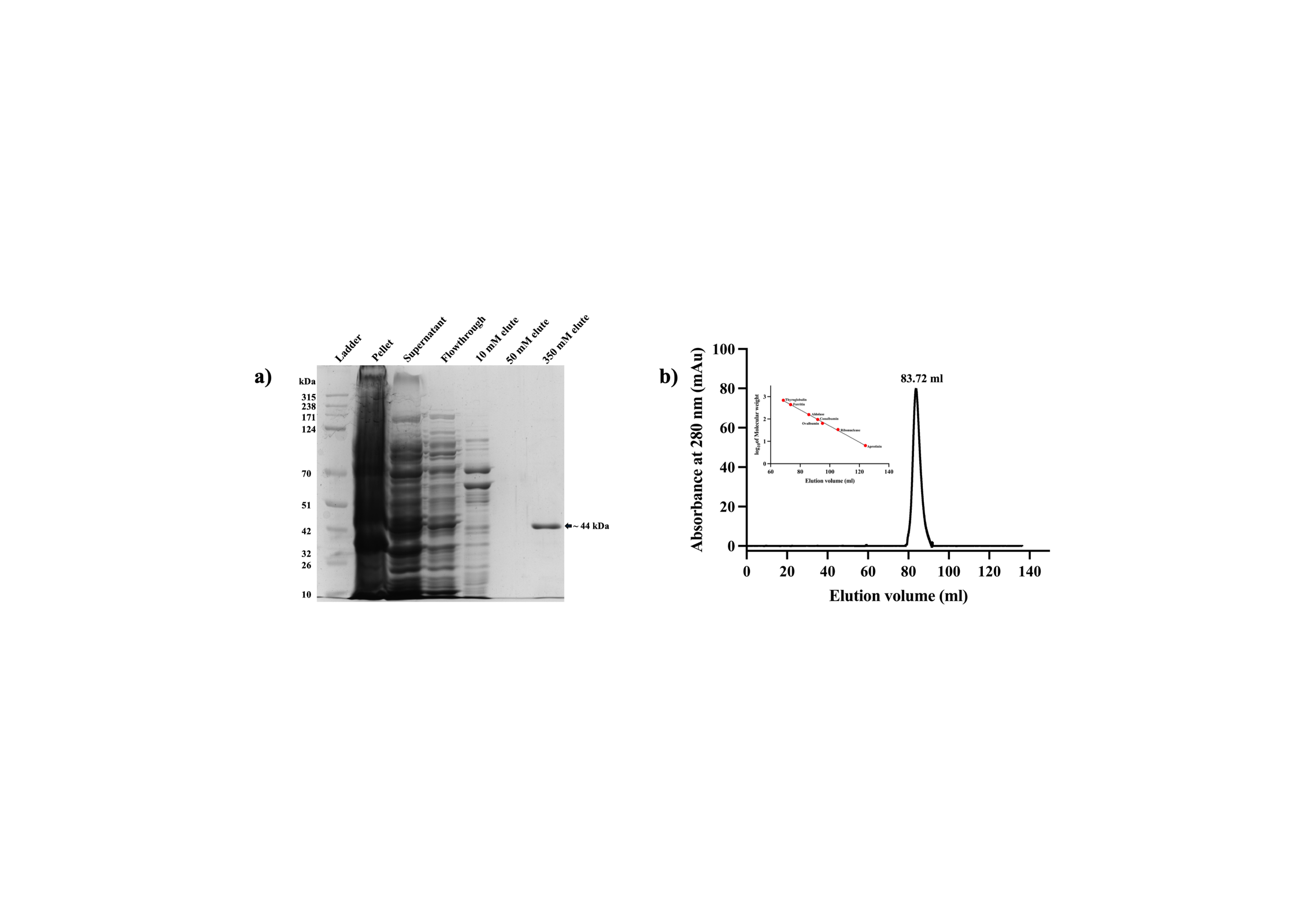

**Figure S3.** The purity of heterologously expressed *C*Las PFP protein was demonstrated by **a)** 12% SDS-PAGE after elution at 350 mM imidazole and **b)** size-exclusion chromatography, which showed a single peak.

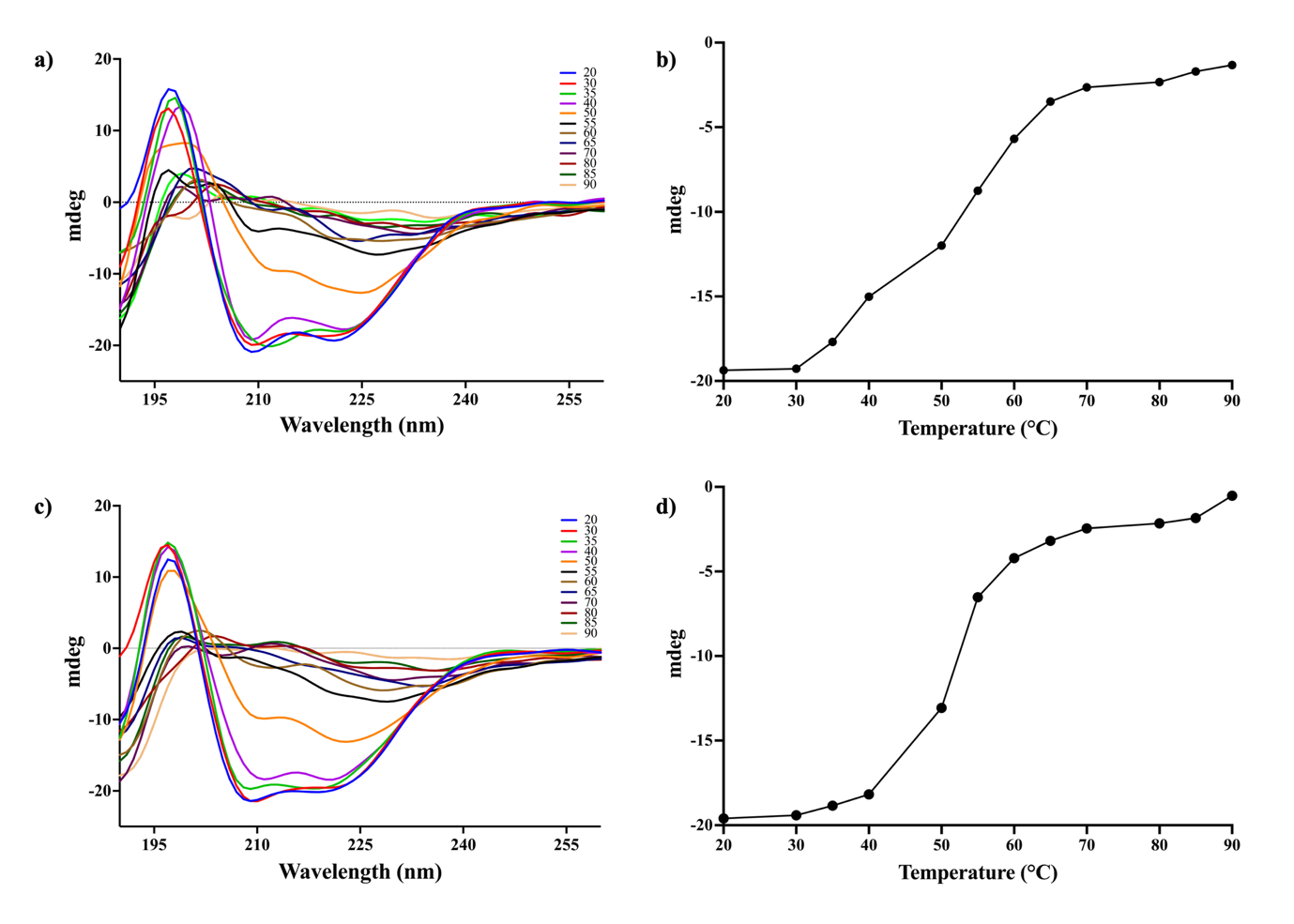

**Figure S4.** CD spectral analysis of *C*Las PFP protein with **a)** F6P and **c)** S7P at different temperatures. The CD spectral signal (mdeg) at 222 nm tracing thermal unfolding of *C*Las PFP with **b)** F6P and **d)** S7P.
